# Simulated visual hallucinations in virtual reality enhance cognitive flexibility

**DOI:** 10.1101/2021.12.14.472556

**Authors:** Clara Rastelli, Antonino Greco, Yoed N. Kenett, Chiara Finocchiaro, Nicola De Pisapia

## Abstract

Historically, psychedelic drugs are known to modulate cognitive flexibility, a central aspect of cognition permitting adaptation to changing environmental demands. Despite proof suggesting phenomenological similarities between artificially-induced and actual psychedelic altered perception, experimental evidence is still lacking about whether the former is also able to modulate cognitive flexibility. To address this, we measure participants’ cognitive flexibility through behavioral tasks after the exposure to virtual reality panoramic videos and their hallucinatory-like counterparts generated by the DeepDream algorithm. Results show that the estimated semantic network has a flexible structure when preceded by altered videos. Crucially, following the simulated psychedelic exposure, individuals also show an attenuated contribution of the automatic process and chaotic dynamics underlying the decision process. This suggests that simulated altered perceptual phenomenology enhances cognitive flexibility, presumably due to a reorganization in the cognitive dynamics that facilitates the exploration of uncommon decision strategies and inhibits automated choices.

## Introduction

Cognitive Flexibility (CF) is commonly defined as the ability to shift attention between competing concepts and alternate behavioral policies to meet rapidly changing environmental demands^1,2^. As the hallmark of creative problem solving, CF enables the generation of original and useful solutions to ill-posed problems^3–5^. As such, CF is considered a fundamental component of cognitive systems, promoting positive life outcomes^6^ and playing a major role in the reduction of aging effects due to cognitive decline^7^. Besides being essential to optimal functioning, many mental disorders, such as psychotic illness^8^ and autism^9^, are marked by a lack of CF. Therefore, elucidating methods underlying the modulation of CF is crucial and valuable to a multitude of fields.

CF is conceived as an emergent property of efficient executive functions, entailing the reconfiguring of one’s behavioral policy to accomplish a new goal. Indeed, CF is optimally supported by the coordinated involvement of several subdomains of executive function, such as working memory and inhibitory control^10^. For instance, higher CF scores have been linked to a higher ability to inhibit automatic responses as measured by the Stroop task^11–13^. As an alternative to executive processes models, associative theories explain CF, and more broadly the creative cognition, as a process through which individuals adapt to new situations through reorganization of their knowledge in the semantic memory space where one stimulus is spontaneously activated by another due to their association^3,4,14,15^.

Across disciplines, researchers have long investigated how to facilitate CF abilities^16–27^. For instance, diversifying experience (i.e., experiencing unusual and unexpected events)^16^, experiencing awe^17^, playing strategy videogames^18^, attending training courses^19^ as well as ingesting psychedelics^20–27^ (e.g., psilocybin, ketamine) resulted in improved CF performance. Critically, among the variety of methods employed, psychedelics have been linked to an enhancement of CF since the middle of the twentieth century^20,28^. At that time, psychedelic drugs were extensively used in experimental research^29,30^, but they were made illegal by the governments of most countries worldwide as a reaction to the counterculture of the 1960s, with the consequence that today there is a lack of compelling evidence regarding their effects. Nowadays, the neuroscientific community has shown a renewed interest in the application of psychedelics to investigate various aspects of brain and cognitive dynamics^25,31,32^, making the application of psychedelic drugs a novel and valuable tool for exploring high cognitive functions, and in particular the flexibility of thought and creativity-related processes^21–25,33^. Indeed, psychedelic experience seems to be associated with an unconstrained mode of cognition, mental imagery, and hyper-associative thinking^25^ that alters the sense of meaning^21,33,34^. Under the influence of lysergic acid diethylamide (LSD), studies found a boosted indirect semantic priming, implying that they may support a broader spread of semantic activation given a stimulus, hence facilitating the recall of distant associations^21,27,31,34^.

Carhart-Harris and Friston^32^ recently provided a theoretical framework that offers an explanation of how psychedelic affects cognitive systems, which in turn may explain for greater CF. According to this model, perception is guided by a predictive coding process that integrates top-down previous beliefs with bottom-up sensory information, thus providing efficient information processing. Although the creative performance may be hampered by prior beliefs eventually skew our view in favor of our prior expectations (i.e., confirmation biases), it was suggested that psychedelic drugs might release such high-level signals and facilitate direct access to conscious experience through the broad communication of bottom-up signals^32^. By expanding the brain’s global flexibility, the subjective experience may become richer and the volume of information, especially mnemonic and sensory, can increase, allowing for new insights to be gained. This phenomenon has been operationalized within the well-known “entropic brain hypothesis”^35^, according to which the subjective experience qualitatively depends on the system’s entropy (i.e., an index of a dynamical system’s disorder).

Although these findings demonstrated the enormous potential of using psychedelics to investigate the neural and cognitive mechanisms of CF, the difficulty of obtaining approval for their use in scientific investigations remains hampered by ethical and legal issues in many countries. To overcome these limitations, Suzuki et al.^36^ proposed a methodology, called the Hallucination Machine, that combines deep convolutional neural networks (CNNs) and panoramic videos, viewed immersively through virtual reality (VR), to simulate biologically plausible “artificial” hallucinations (animals, faces, etc.). Using behavioral measures, they found that this simulation induced visual perceptual phenomenology qualitatively similar to psychedelics^36^. Moreover, Greco et al. found similar brain patterns between psychedelic drugs and the artificially-induced altered perceptual phenomenology, with an increased entropic brain dynamics and global functional connectivity^37^. These findings further support such methodology the study of the phenomenological aspects of the psychedelic experience.

Despite earlier scientific studies demonstrating a beneficial effect of psychedelics on CF, experimental evidence is lacking whether the Hallucination Machine might modulate CF similarly to the actual psychedelic experience. Since DeepDream altered perceptual phenomenology in VR appears to be qualitatively comparable to psychedelic experience^36,37^, and the latter seems to modulate CF^21–24,33^, it is noteworthy to examine the effects of DeepDream altered perceptual phenomenology in VR on CF. Importantly, if artificially-induced perceptual phenomenology is able to mediate changes in CF, it could be used as a potentially novel, ecological, and controlled tool to investigate CF, as well as the underlying neural mechanisms.

In the present study, we exposed participants to two series of video clips, one depicting regular natural scenes and one modified by DeepDream^38^, the same algorithm implemented by Suzuki et al.^36^ to generate artificial visual hallucinations. After the exposure to each of the video sessions in VR, participants completed both the Alternative Use Task (AUT)^39^ and a mouse-tracking version of the Stroop task^40,41^ for assessing their CF, and finally, a brief version of the altered states of consciousness (ASC) questionnaire^42^ to assess their phenomenological experience. We tested whether DeepDream-induced altered perceptual phenomenology enhanced CF as compared to regular perceptual phenomenology. To achieve this goal, we used a within-subject design and a combination of computational techniques such as network science for the estimation of semantic networks from the AUT responses^4,15^, and the drift-diffusion conflict model (DCM)^43–45^ for modeling the accuracies and reaction times from the Stroop data.

In the context of the AUT, CF is usually measured by the quantity of switches between semantic categories taps from participants. The flexibility of the AUT responses is usually measured by a panel of judges, bringing with it all the issues related to subjectivity and therefore the replicability of results. Therefore, here we adopt a method recently developed based on network science methodology and percolation theory to examine CF^4,15,46^, yet extensively used to investigate how semantic memory organization may aid flexible thinking^14,47^. Thus, characterized by higher connectivity and shorter overall distances between concepts, the semantic network allows for more efficient spreading of activation processes throughout the semantic space, which may contribute to the generation of more distinctive ideas^4,15^. Regarding the Stroop task, CF is quantified as the ability to inhibit automatic responses^11–13^. Here, we modeled Stroop data using the DCM which enables us to explain decisions in conflict situations in terms of cognitive control and spontaneous processing mechanisms^43–45^. We additionally examined the mouse trajectories during the Stroop task in order to quantify the level of tortuosity elicited by the altered perceptual phenomenology and control condition.

Motivated by recent studies suggesting a potential effect of psychedelics on the increase of brain’s global flexibility^24,31,32^ and precisely on the spread semantic activation^21,33,34^, we expected participants to exhibit a more flexible structure of the semantic network after the altered perceptual phenomenology condition compared to the control condition, as a result of condition-related differences in these search processes. We further tested this hypothesis by examining the robustness of the semantic networks under targeted attacks using percolation analysis, assuming that the higher the robustness of a semantic network, the higher its flexibility^4,46^. Moreover, we predicted an attenuated contribution of prior knowledge to the participants’ decision-making and increment in the efficiency of the performance as a result of experimental stimulation^32^. Indeed, the inhibition of automatic responses is an important characteristic of CF^12,13,48^. Finally, a more chaotic pattern of the mouse trajectories in the experimental condition would signify that the perturbation at the participant’s lower perceptual level expectations ultimately affected higher-level cognitive processes^35^.

## Results

Data were collected from 52 individuals in an equipped VR lab. Participants were exposed to the series of original videos (OR condition, Fig. 1a) followed by DeepDream videos (DD condition, Fig. 1a) in VR. The order of conditions was counterbalanced across participants. Immediately after the videos’ presentation in VR in each condition, volunteers performed two behavioral tasks and a questionnaire on a computer screen, always in the same order (Fig. 1b). The first task was the AUT^39,49^ (Fig. 1c), in which participants were asked to list as many unusual uses as they could think of to four cue words (e.g., “Newspaper”). The second task was a mouse-tracking version of the Stroop task^40,41^ (Fig. 1d), requiring participants to click on the button which represented the color of the target word while ignoring its meaning. After the tasks, the ASC^42^ questionnaire was administered to measure specific dimensions of the participants’ subjective experience. The ASC was used to probe the effectiveness of the DD videos to simulate visual hallucinations, following Suzuki et al.^36^. We performed a two-tailed paired permutation *t*-test (α = 0.05, 10000 iterations) and Cohen’s d (*d*) measure of effect size to compare responses to the ASC items following the two conditions. Results showed a significant increase of the ratings in DD compared to OR (Fig. 1e, see also Table 1 in Supplementary Materials) on the following dimensions: vivid (*p* = .016, *d* = 0.45), patterns (*p*= <.001, *d* = 1.79), imagery (*p* = <.001, *d* = 2.63), intensity (*p* =.005, *d* = 0.46), strange (*p* = <.001, *d* = 2.85), space (*p* = <.001, *d* = 0.81), muddle (*p* = <.001, *d* = 1.03), and spirit (*p* = .013, *d* = 0.50).

**Fig. 1.**
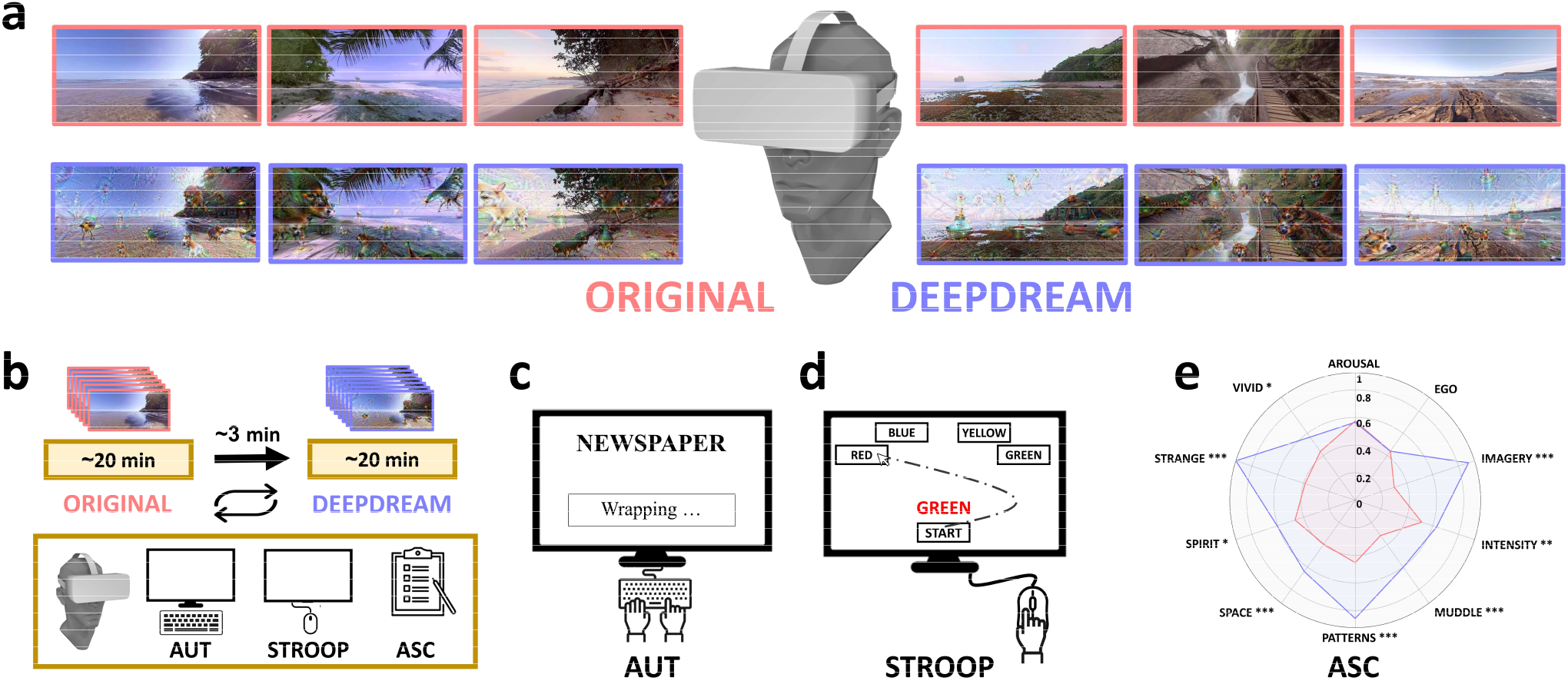
Experimental design and stimuli. **(a)** Visual stimuli presented in VR. They were panoramic 360° videos depicting natural scenes (red frames) and their DeepDream modified counterparts (blue frames). **(b)** Experimental design. Recurring arrows refer to the counterbalanced order of the conditions across participants. **(c)** Schematic example of the AUT. **(d)** Schematic example of the Stroop task. **(e)** Radar plot of the ASC results. Red and blue areas represent OR and DD conditions, respectively. Statistical significance. * -*p* < .05; ** - *p* < .01, *** - *p* < .001.

**Table 1.**
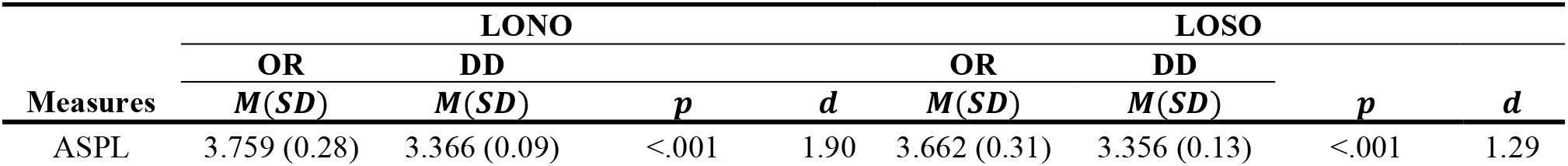
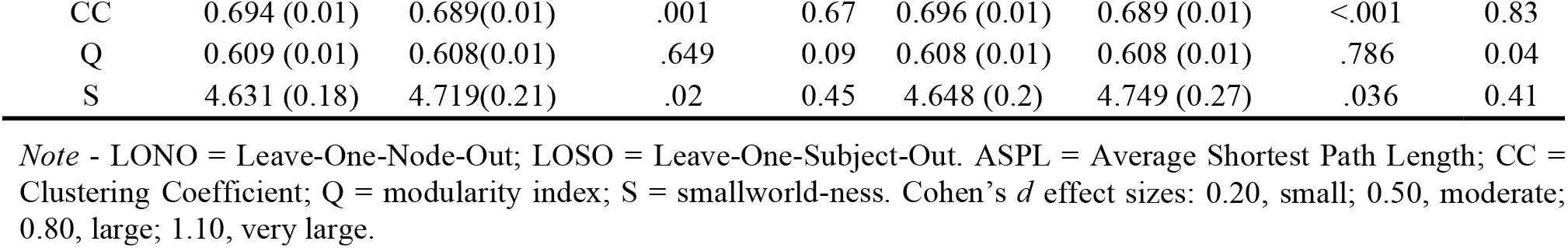
Results from the paired two-tailed permutation t-test of the partial networks comparing the OR and the DD conditions

### Semantic network structure

The AUT responses were pre-processed and resulted in 567 unique responses in total across the sample. The McNemar’s chi-squared test and the Phi measure of effect size (φ) were used to examine whether there was a difference in the proportion of unique responses between the condition. In the OR condition, participants generated 339 of the total unique responses (196 of which were not given in DD), and in the DD condition, participants generated 371 of these responses (228 of which were not given in OR). The proportion of the number of unique responses in DD (65.5%) was larger than the proportion in OR (59.8%), even though this effect was not significant (*χ*^2^(1) = 2.266, p = .132, φ = 0.06). After this step, we constructed a group-level semantic network for each condition using the preprocessed AUT unique responses. We adopted a recent network science approach^15,50^, which allows examining the structure of a semantic memory network, depending on how often the responses co-occur across the sample. According to the spreading activation model^51^, a concept in semantic memory is represented as a node of a semantic network, and it is connected to other nodes based on a semantic similarity principle. Thus, the essential principle of the adopted method^15,50^, is that semantically related concepts tend to be located close together in the semantic memory and have stronger connections. In order to construct the semantic network, we first selected only the unique AUT responses that matched between conditions and were collected from at least two participants, which resulted in 63 responses (nodes). Then, we computed the cosine similarity between the binary vectors associated with each selected unique response, representing which participant made that response, in a pairwise fashion. This resulted in an undirected weighted semantic network for each condition, with the unique response as nodes and the cosine similarity as links. After filtering^52^ the semantic networks, their organization was analyzed using the following network measures, commonly examined in semantic network research: Clustering Coefficient (CC)^53^, Average Shortest Path Length (ASPL), modularity index (Q)^54^, and Small-worldness measure (S)^55^. The CC is a quantifier of network connectivity, measuring the degree to which nodes are near-neighbors and tend to cluster together in the network. A higher level of CC indicates a better local organization, as well as a more interconnected network^53^. The ASPL also characterizes the strength of the association between nodes, indicating the averaged shorter steps between any two pairs of nodes. As such, the ASPL has been considered a lead factor in the spread of the activation^14,56^; the more the ASPL is reduced the higher the chances of reaching a wider number of connections. Finally, the Q allows us to quantify the modularity of the network, i.e. ways in which a network is divided into sub-networks^54^. Finally, the S measure is a measure of how efficient, flexible, and chaotic a network is^55^. Characterized by high local connectivity (CC) and short global distances between nodes (ASPL), the S measure contributes to an optimal diffusion search and retrieval through semantic memory.

Results from the comparison of the full network revealed qualitative (Fig. 2a) and quantitative (Fig. 2b-c) differences between the OR and DD networks’ structures. The OR network appeared to be more spread out than the DD network (Fig. 2a). Conversely, the DD network showed a reduced distance between nodes, as reflected in the lower ASPL (Fig. 2b). The semantic network of the DD condition showed lower structural (ASPL = 3.388, Q = 0.607) and higher flexible (S = 4.678) values compared to the network of the OR condition (ASPL = 4.052, Q = 0.614, S = 4.565). The CC measure showed a small difference, with a higher value in the OR network (CC = .6906) than the DD network (CC = .6905). We statistically examined the validity of our findings by applying two complementary approaches, the Leave-One-Node-Out (LONO) and the Leave-One-Subject-Out (LOSO). In the LONO procedure, we iteratively computed the topological quantifiers on the partial networks resulting from the exclusion of one node at each iteration, for every node. In the LOSO procedure, for each iteration, we excluded one participant and repeat the pipeline for building the semantic networks and computing the network measures, for every participant. A two-tailed paired-samples permutation t-test (α = 0.05, 10000 iterations) were computed on each measure for comparing the conditions in both the LONO and LOSO procedures. Results showed that DD had a significantly smaller ASPL, CC, and higher S compared to OR, confirming the analyses on the full networks, with effect sizes ranging from moderate to very large (Fig. 2c). The Q parameter did not reach a statistically significant difference between conditions. For statistics see also Table 1.

**Fig. 2.**
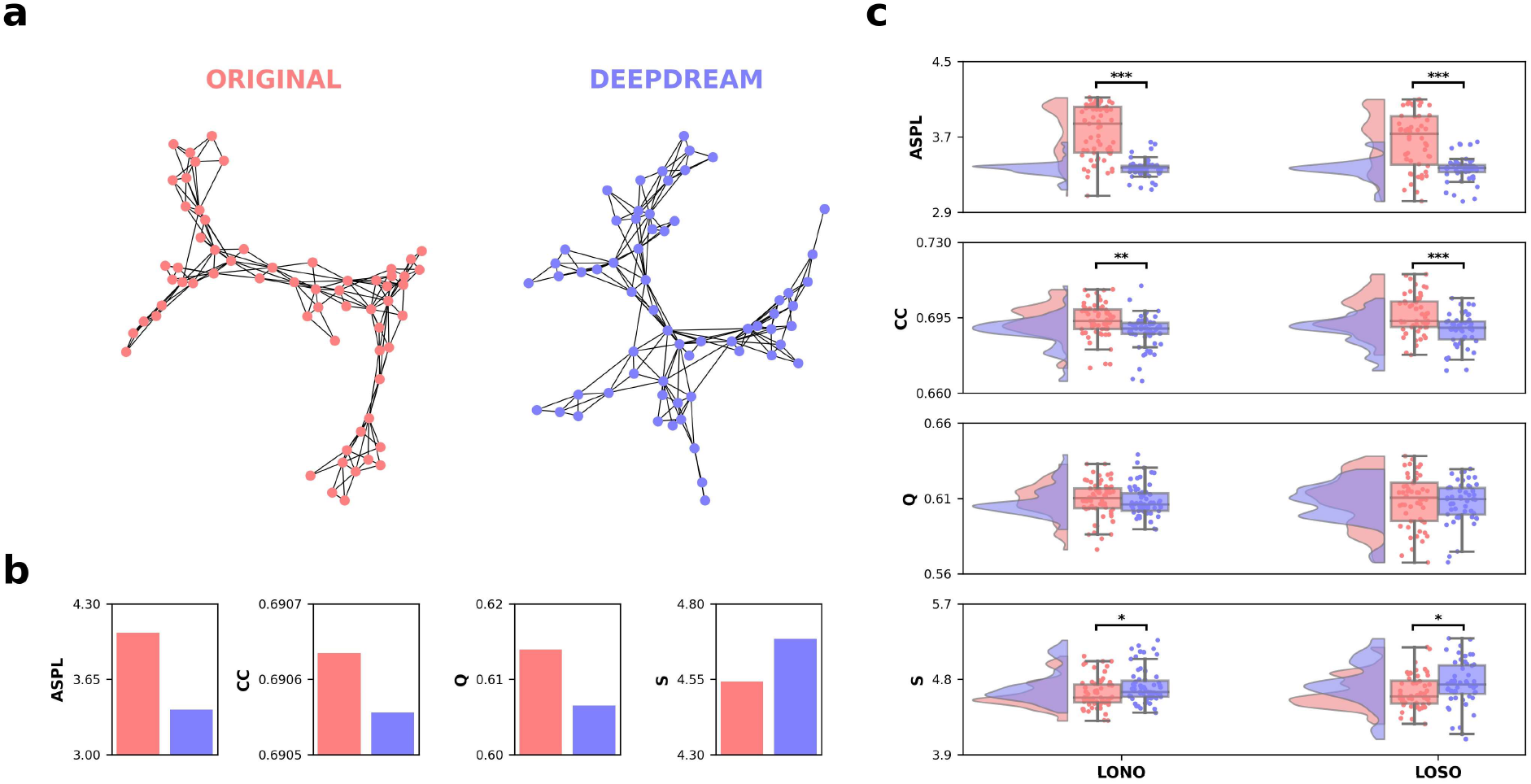
Semantic networks, topological quantifiers and statistical results. **(a)** Undirected, unweighted semantic networks of the OR and DD conditions, visualized using the spring layout, with nodes as unique AUT responses and edges as cosine similarity. **(b)** Barplots depicting the topological quantifiers of the full networks. **(c)** Raincloud plots represent the results from the LONO and LOSO procedures on the topological quantifiers. Horizontal black bars represent statistical significance.

### Network Percolation

We also assessed the robustness of the semantic networks using a network percolation analysis approach^4^ to probe the resiliency of the network under targeted attacks. Here, the weighted networks previously constructed from the AUT data were used. In the percolation analysis, networks are “attacked” by removing links with weight strength below an increasing threshold, called the percolation step. In each percolation step, we measure the size of the Largest Connected Component (LCCS), which is the largest cluster of nodes connected only to each other. Once the percolation process reached its end, we computed the percolation integral (*ϕ*), which is the area under the curve representing the LCCS across the percolation steps. We applied this analysis on the full networks of both conditions, finding that the percolation integral of DD (*ϕ* = 22.15) was larger with respect to OR (*ϕ* = 17.15), meaning that the OR network broke apart faster compared to the DD network, as illustrated in Fig. 3a. In Fig. 3c, we illustrated how the networks appeared throughout the percolation process and can be appreciated the difference in the LCCS between the conditions at different percolation steps, denoting the DD network’s robustness compared to the OR network. To determine the statistical significance of our findings, we implemented three approaches: LOSO, LONO, and the Link Shuffling analysis (LS). As mentioned above, when computing the LONO and LOSO procedures, we iteratively excluded each node or participant, ran the percolation process on the resulted networks, and computed *ϕ* . For the LS analysis, we randomly exchanged pairs of links (~1700) in the network and computed *ϕ*, repeating this procedure for 500 iterations. A two-tailed paired-samples permutation t-test (α = 0.05, 10000 iterations) was computed on *ϕ* for comparing the conditions in the LONO, LOSO, and LS procedures. Overall, these analyses revealed similar results (Fig. 3b), namely that the average *ϕ* of DD was significantly larger than OR (all *p* < .001) and very large effect size (*d*_*LONO*_= 7.01, *d*_*LOSO*_= 1.89, *d*_*LS*_ =5.08; for statistics see also Table 2).

**Fig. 3.**
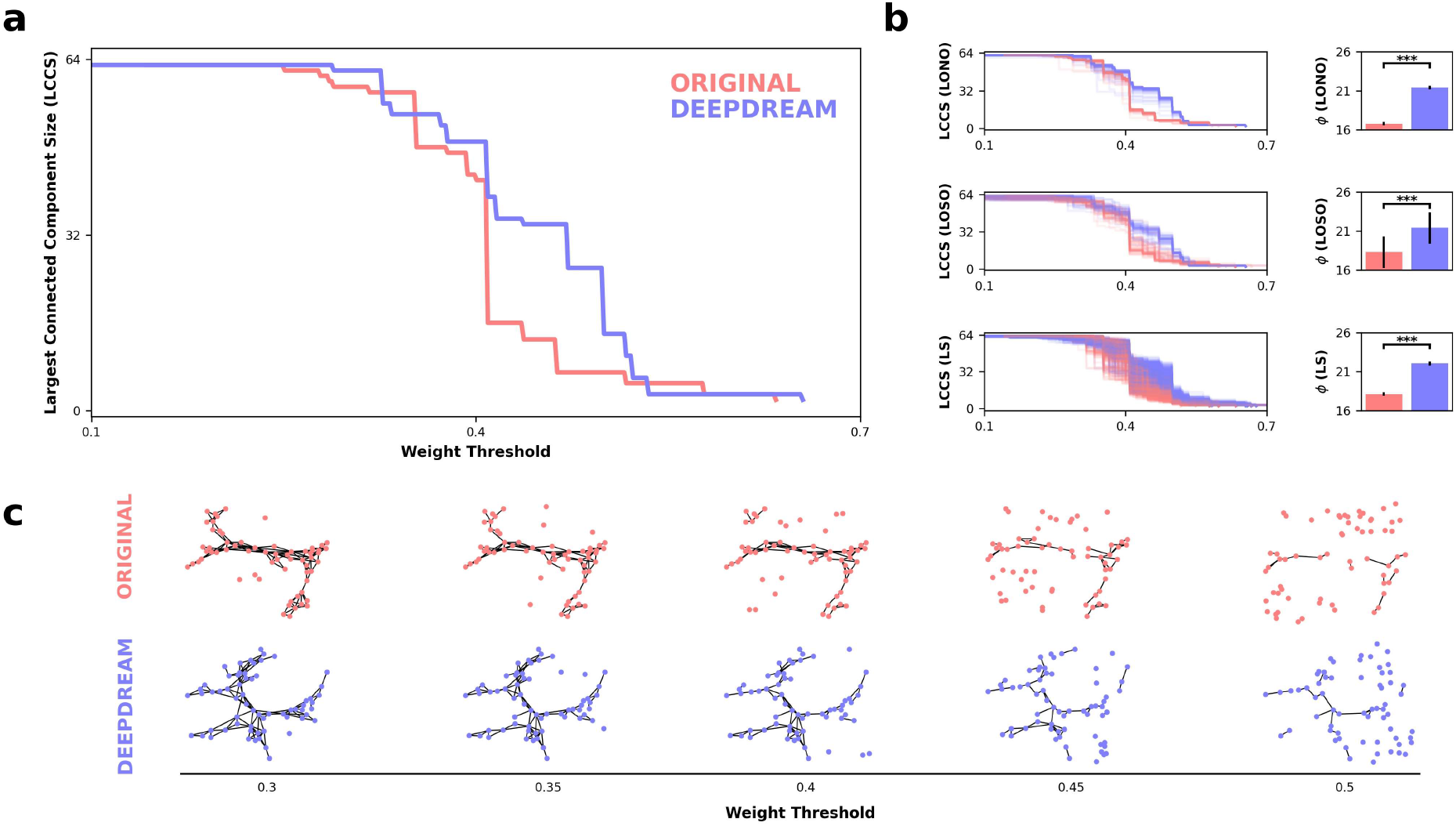
Network percolation and statistical results. **(a)** Line plot representing the percolation process of the OR (red) and DD (blue) full networks. The x-axis represents the weight threshold, starting from the smallest weight in the network (0.1) to a weight strength in which the giant component is smaller than three nodes (0.7). **(b)** On the left, line plots of the LONO, LOSO, and LS procedures. Each line is an iteration, colors encode the conditions. On the right, barplots show the *ϕ* between conditions and across the three procedures. Error bars represent the standard error of the mean (SEM). Horizontal black bars represent statistical significance. **(c)** OR and DD semantic networks undergoing the percolation process, visualized at different weight thresholds.

**Table 2.** Results from the two-tailed paired-sample permutation *t*-test on percolation integral comparing the OR and the DD conditions.

| Method | OR | DD | $p$ | $d$ |
| --- | --- | --- | --- | --- |
| Leave One Node Out | 16.75 (0.27) | 21.4 (0.89) | < .001 | 7.01 |
| Leave One Subject Out | 18.31 (2.07) | 21.42 (1.02) | < .001 | 1.89 |
| Link Shuffling | 18.13 (0.73) | 22.06 (0.81) | < .001 | 5.08 |

*Note* - Mean and standard deviation in parenthesis. Cohen's $d$ effect sizes: 0.20, small; 0.50, moderate; 0.80, large; 1.10, very large.

### Drift Diffusion Conflict Modelling

Stroop data were pre-processed by removing outliers both in terms of reaction times (RT) and accuracies. After this step, we compared accuracy and RT data across Stroop conditions using a paired-samples two-tail permutation *t*-test (α = 0.05, 10000 iterations, Fig. 4a). We did not find any significant difference between OR and DD in the accuracy of congruent (*p* = .787, *d* = 0.10) and incongruent trials (*p* = .098, *d* = 0.20), neither in the RT of congruent (*p* = .749, *d* = 0.03) and incongruent trials (*p* = .642, *d* = 0.04). Then, we fitted the DCM to the RT and accuracy data for each participant and condition separately. The DCM is a computational model for conflict tasks such as the Stroop^45^, modeling the decision-making of participants as a superimposition of a controlled and automatic process. The model assumes that the automatic process increases the rate of evidence accumulation (although based on task-irrelevant information) in the congruent trials while decreasing it in the incongruent trials (Fig. 4b). Model fitting was performed by simulating trials from the model and matching the cumulative distribution function (CDF) of the RT data and the conditional accuracy function (CAF) using the Nelder-Mead optimizer (Fig. 4c, see methods for details). We compared 4 parameters from the model fitting (Fig. 4d), namely the amplitude of the automatic process (*α*), the decay of the automatic process (*τ*), the drift of the controlled process (*δ*) and the decision boundary (*β*). Statistical significance was assessed with two-tailed paired-samples permutation *t*-test (α = 0.05, 10000 iterations). We found that *α* was significantly reduced in DD compared to OR (*p* = .012, *d* = 0.51), indicating that the contribution of the automatic process to the participants’ decision-making was reduced in DD. We did not observe any difference in *τ* (*p* = .986, *d* = 0.01), while a higher average value, although not statistically significant, of *δ* (*p* = .282, *d* = 0.16) and *β* (*p* = .507, *d* = 0.11) was observed for OR with respect to DD.

**Fig. 4.**
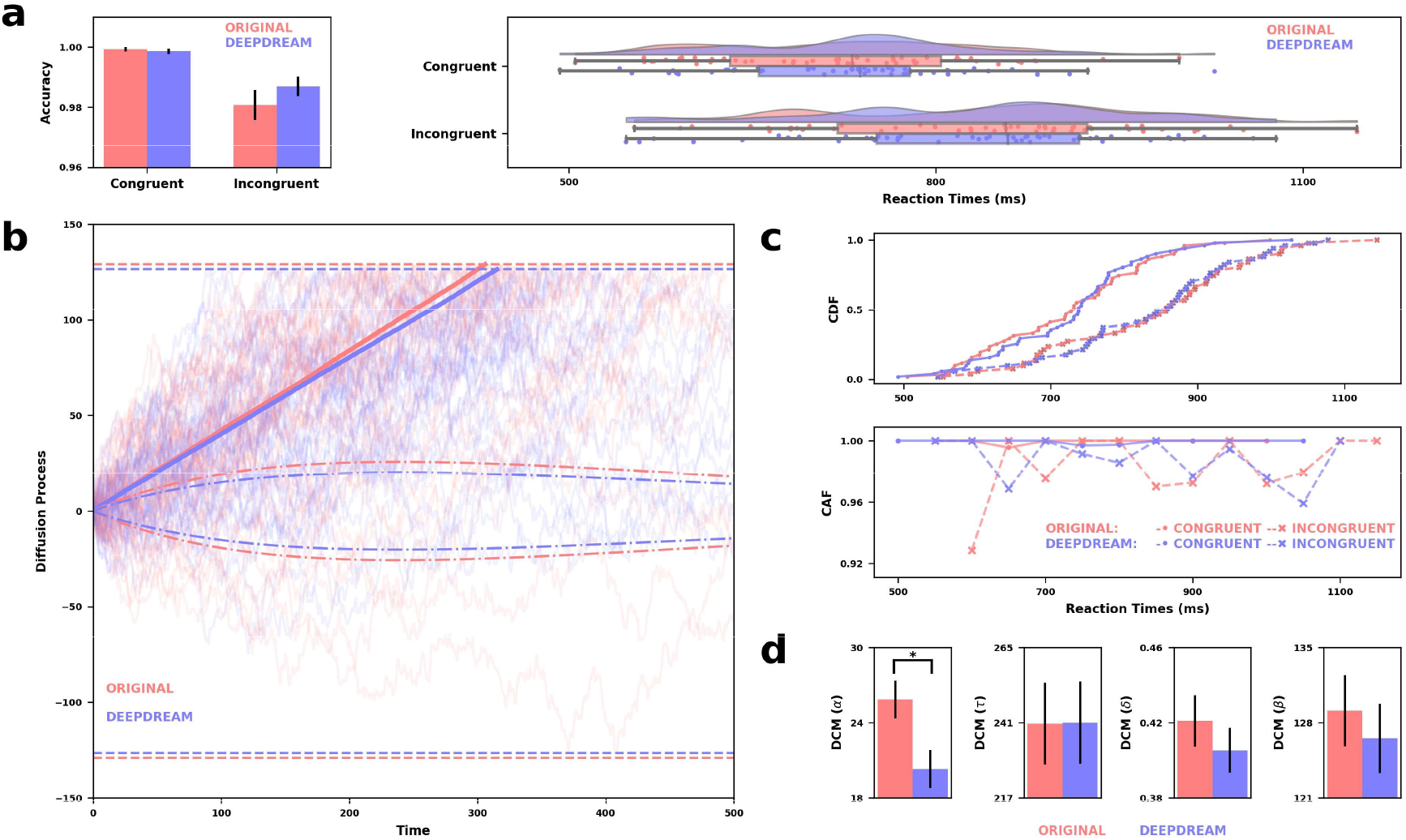
**(a)** On the left, bar plots of the accuracies between OR (red) and DD (blue) conditions, and congruent and incongruent trials, with error bars indicating SEM and horizontal bars representing statistical significance. On the right, raincloud plots of the RT across Stroop conditions. **(b)** Graphical depiction of the results from the DCM. Semi-transparent solid lines are 10000 trials simulated per condition using recovered parameters from model fitting. Solid lines are the average of the simulated trials per condition. Dash-dotted lines represent the automatic process, while dashed lines represent boundaries. **(c)** Group-level statistics were used for fitting the DCM. On the top, the cumulative distribution functions (CDF), while on the bottom are the conditional accuracy functions (CAF), both plotted across Stroop conditions. **(d)** Barplots indicating the estimated parameters from DCM. Error bars indicating SEM and horizontal bars representing statistical significance.

### Mouse trajectory analysis

We analyzed the Stroop data also in terms of mouse trajectories. Trajectories were firstly selected based on the exclusion criteria from RT and accuracy data and including only the correct trials. We extracted from the raw trajectories only the data points in which the cursor actually moved. Then, we time normalized the trajectories using linear interpolation, making all trajectories with the same number of data points (n = 101), and spatially aligned them in order to have the same initial and ending point. Finally, we computed the area under the curve (AUC) between the trajectories and optimal path, the permutation entropy (PE), and the number of deviations (D) along the x and y coordinates and the Euclidean distance (ED) and the velocity (V). Statistical significance was assessed with a two-tailed paired-samples permutation *t*-test (α = 0.05, 10000 iterations). We found that trajectories were closer to the optimal path in the congruent trials with respect to the incongruent ones in both OR and DD conditions (Fig. 5a). We also found that AUC was slightly lower in OR compared to DD, meaning that OR trajectories were more efficient, even though this result was not significant in both congruent (*p* = .555, *d* = 0.04) and incongruent (*p* = .839, *d* = 0.01) trials. Moreover, we also found that PE was significantly higher in DD compared to OR on the incongruent trials along both the x (*p* = .045, *d* = 0.24) and y (*p* = .048, *d* = 0.20) coordinates. No significant difference was observed on the congruent trials in both the x (*p* = .485, *d* = 0.07) and y (*p* = .542, *d* = 0.06). Similarly, we found that D was significantly higher in DD compared to OR on the incongruent trials along the y (*p* = .042, *d* = 0.24) coordinate but not x (*p* = .141, *d* = 0.22). No significant difference was observed on the congruent trials in both the x (*p* = .231, *d* = 0.16) and y (*p* = .858, *d* = 0.02). Also, we did not observe any significant difference in ED (congruent *p* = .685, *d* = 0.04; incongruent *p* = .621, *d* = 0.07) nor in V (congruent *p* = .844, *d* = 0.02; incongruent *p* = .781, *d* = 0.03). Surprisingly, we observed that PE, ED, and D were generally higher in congruent trials with respect to incongruent trials in both conditions. Furthermore, we used Gaussian Mixture Models (GMM) to estimate macro-states’ trajectories in order to better characterize the decision process of participants. Model selection evidenced that a GMM with 4 clusters was the best fitting model. We decided to label these clusters as Initiation, Prediction, Evaluation, and Termination states. As shown in Fig. 5c, Initiation and Termination states pertain to the starting and ending phase of the trajectories, respectively. The Prediction state was subsequent to the Initiation and was considered as a moment in which participants made their first guess about the correct outcome of the trial. After this, the Evaluation state is a phase in which participants could in principle change their first prediction and be attracted more towards other targets. We computed the transition matrices among these states between conditions and split between congruent and incongruent trials (Fig. 5d). Boradly, we found, as expected, that it was more probable to remain in a certain state with respect to switch since these states can be conceived as mostly sequential states of the decision process. Finally, we computed the Dwell Time (DT), defined as the average lifetime of a state (Fig. 5e). We found a significant difference in the termination state between OR and DD in congruent (higher OR, *p* = .019, *d* = 0.22) but not in incongruent trials (*p* = .187, *d* = 0.12). Also, in the Prediction state DD had a significant higher DT compared to OR in congruent trials (*p* = .041, *d* = 0.19), while no difference was observed in incongruent trials (*p* = .210, *d* = .08). No significant difference was observed in Initiation (congruent: *p* = .582, *d* = 0.04; incongruent: *p* = .145, *d* = 0.12) and Evaluation states (congruent: *p* = .757, *d* = 0.02; incongruent: *p* = .744, *d* = 0.02).

**Fig. 5.**
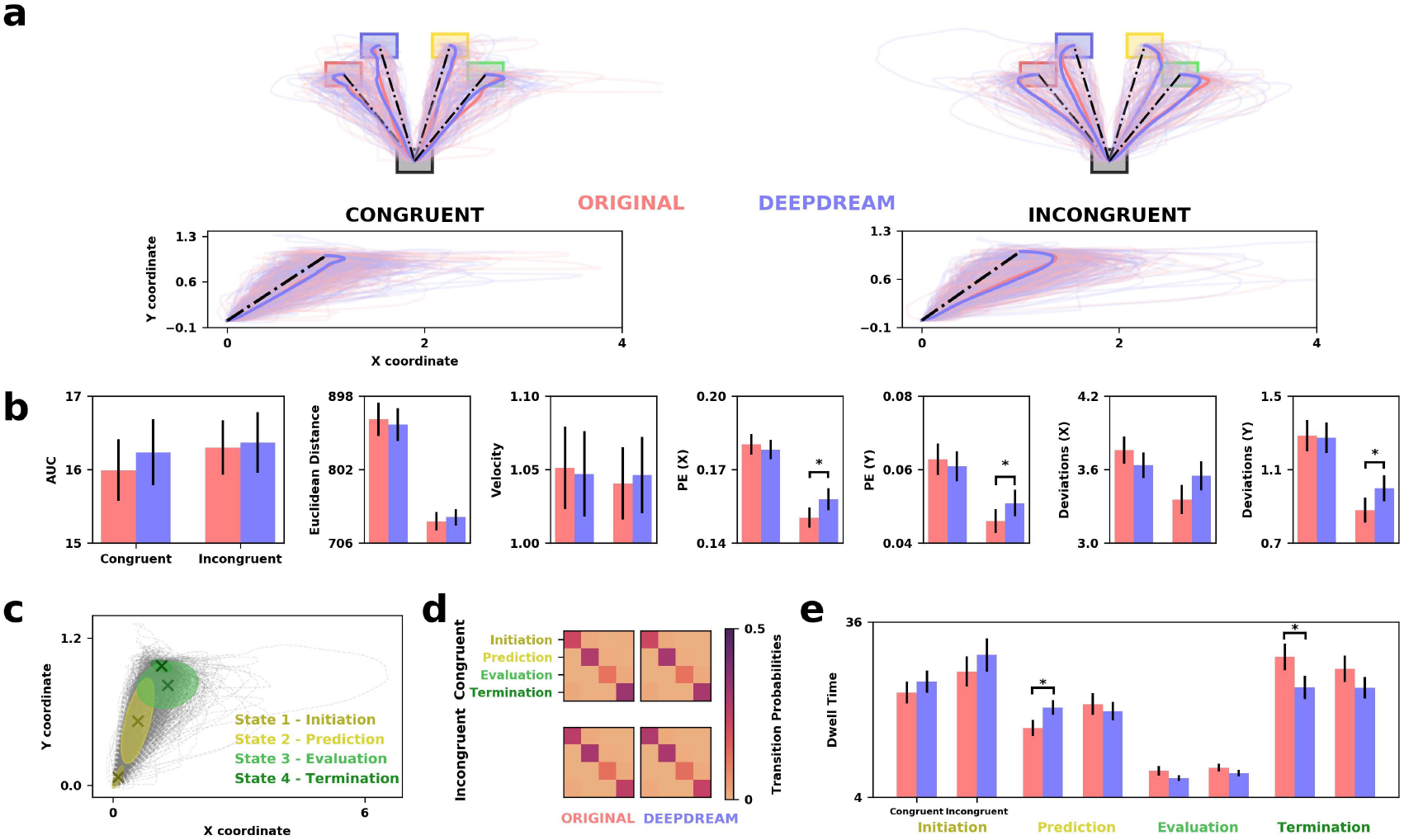
**(a)** On the top, mouse trajectories of OR (red) and DD (blue) spatially aligned to the four targets and divided by congruent (left) and incongruent (right) trials. On the bottom, mouse trajectories spatially aligned to the same initial (0,0) and ending (1,1) point. **(b)** Barplots depicting the measures applied to the mouse trajectories. Error bars indicating SEM and horizontal bars representing statistical significance. **(c)** GMM clusters of mouse trajectories. Grey dashed lines represents the trajectories, crosses are the centroids of each cluster while ellipsoids are the covariances. **(d)** Transition probability matrices for each condition and split between congruent and incongruent trials. **(e)** Barplots representing dwell time for each state. Error bars indicating SEM and horizontal bars representing statistical significance.

## Discussion

Despite scientific studies demonstrating a beneficial effect of psychedelics on CF^20–26^, the effect of artificially induced altered perception on CF has remained largely unexplored. In the present study, we applied the DeepDream algorithm to a series of panoramic videos of natural scenes with the intent to simulate visual hallucinations^36^. Then, we tested whether DeepDream enhanced CF as compared to regular perceptual phenomenology. To achieve this goal, we used a within-subject design in which participants were exposed to DD and OR (control) video sessions in VR. After the video presentation, CF was assessed by both the AUT^39^ and a mouse-tracking version of the Stroop task^40,41^, while participants’ phenomenological experience was measured through a brief version of the ASC questionnaire^42^. ASC revealed a significant increase in DD over OR of perceptual (‘patterns’, ‘space’) and imaginative dimensions (‘imagery’, ‘strange’, ‘vivid’, ‘muddle’) as well as the overall intensity and mystical quality (‘spirit’) of the experience. Our results are largely in line with the findings reported by Suzuki et al. using the Hallucination Machine, as well as studies reporting alteration in participants’ subjective experience after pharmacological administration of psilocybin^36^. Importantly, our findings corroborate the effectiveness of the DeepDream stimulation procedure on the modulation of distinct aspects of altered states of consciousness, especially visual hallucinations, avoiding the extensive systemic effects produced by pharmacological interventions.

Next, we applied a network science methodology in order to construct cognitive networks from the AUT responses. Within this framework, we estimated and compared the properties of the semantic network of each condition^15^ and performed a percolation analysis on these networks^4^. Here, we assumed that a more smallworlded network should facilitate semantic search processes by connecting weakly related concepts, hence enhancing CF. We found that DeepDream exposure significantly influenced the network organization, leading to a reduction of the shortest path between nodes and an increase S, with respect to the OR condition. Consistent with our findings, a higher value of S in the semantic has been previously associated with high creative individuals^15,47,57^. Yet, in our results we found significantly higher connectivity in the OR over DD, suggesting that the reduced ASPL, in DD over OR, was the driving effect of these results, augmenting the chances of reaching a wider number of semantic connections^56,58^. We interpret these findings as showing that the DD network had a more efficient and flexible structure as compared to the OR network, indicating a higher level of CF in the participants elicited by DD^14^. A similar effect was also found by^21,33,34^ suggesting that LSD and related psychedelics increase the spread of semantic activation. Critically, the percolation analysis yielded consistent results, indicating that the cognitive network of the DD condition was significantly more robust to the percolation process, as exhibited by a higher percolation integral (i.e., DD network breaks apart slower than OR). The cognitive network outcomes are further consistent with current theories of semantic memory described as a dynamic system, able to change its organization in the short-term period^21,59–61^. Therefore, our findings may also provide support for process-based change in the conceptual networks.

Furthermore, Stroop data were firstly analyzed in terms of accuracy and RT by means of an integrated computational framework, namely the drift diffusion model^43,44^. We implemented a specific version of this model, the DCM, tailored for conflict tasks^45^. We found that, despite the drift parameter *δ* of the controlled process was similar between condition, DD had a significantly lower amplitude of the automatic process compared to OR, suggesting that automatic processes contributed less to the overall decision process in DD. Indeed, the inhibition of automatic responses is an important characteristic of CF^12,13,48^, therefore we interpreted these results as a confirmation of our previous network findings but from a different perspective, i.e. inhibitory control. Contrary to our expectations, we did not observe an increment in the efficiency of the performance due to DD. Although we did not find any significant results, we can speculate that an explanation for this observation was that the *δ* parameter was higher in OR, but compensated by a lower *β* parameter in DD. Also, this might be due to the fact that DD modulated not only CF, but also other aspects of cognition that might have interfered with the performance on the Stroop task.

These preliminary interpretations were also corroborated by the analyses performed on the mouse trajectories, evidencing general differences in the strategies adopted by participants to reach the target between conditions. Firstly, we found a similar pattern of general performance with respect to the DCM results, i.e., OR had a lower AUC compared to DD, albeit not significant as in the DCM. Importantly, we found that the trajectories in the incongruent trials were characterized by a higher level of tortuosity (higher PE and D) in DD compared to OR. Also, GMM clustering analysis revealed a higher tendency to stay in the early stages of the decision process in DD with respect to OR in congruent trials (higher Dwell Time of DD in the Prediction state and of OR in the Termination state). These findings clearly show how DD affected pervasively participants’ cognitive abilities by perturbing their lower-level perceptual expectations that ultimately affected higher-level cognitive processes.

Crucially, these results seem to corroborate at the behavioral level the findings from Greco et al.^37^, in which they observed a general increased entropic brain dynamics due to DeepDream exposure. Therefore, it seems reasonable to interpret these mouse trajectories findings in light of the entropic brain hypothesis^32,35^, according to which, during psychedelic experiences, the brain is supposed to operate at a criticality state, where it has access to a larger repertoire of physical states and therefore has a more chaotic dynamics. Here, we speculate that the tortuosity of the trajectories alongside the increased latency in resolving the uncertainty of the choice at the behavioral level might be explained by the higher entropic dynamics at the neural level. As stated above, these explanations could also be in favor of the observation that DD did not improve participants’ performance, since it is more difficult to reach a higher level of efficacy with such a chaotic regime in the brain dynamics.

Overall, our findings can also be interpreted in line with recent evidence suggesting that diversifying experience, loosely defined as highly unusual and unexpected events, can lead to an enhancement of CF ^16,62^. Our contribution to this line of research was to provide a quantitative and parametric approach to experience diversification by employing the DeepDream stimulation procedure.

Some limitations to our study exist. First, due to time limitations switching tasks were not administered. An interesting future direction could be the extension of CF measures to adopt in order to better characterize how DeepDream affects CF, conjunctly with other executive functions. Second, although the AUT is a valuable task to study creativity, it is usually administered as an open-ended task, whereas we required participants to only generate single-word responses. This requirement potentially constrains participants’ responses in this task. Thus, future research is needed to replicate and extend our findings in more standard tasks used to assess semantic memory networks, such as free associations and semantic fluency tasks^56^. Third, we analyzed the semantic networks at the group level, aggregating across individuals and thus ignoring unique individual differences. Follow-up research should replicate our findings using the DeepDream manipulation, by estimating individual-based semantic networks.^63^. Lastly, future studies are encouraged to augment the stimulus set by exploring different parameter settings of DeepDream, for instance by generating different videos that match, from low-level to high-level features, their original counterparts; this would clarify how and to what extent the low-level features modulate CF.

In conclusion, our findings provide evidence that simulated altered perceptual phenomenology enhances CF, presumably due to a reorganization in the cognitive dynamics that facilitates the exploration of uncommon decision strategies and inhibits the prevalence of automatic choices. We also showed how the use of recent deep learning models could furnish cognitive science research with a new tool to investigate low- and high-level cognition. Our study illustrates the strength of applying DeepDream-induced altered in studying cognitive processes, as well as further investigations on similar techniques for fostering cognitive flexibility.

## Methods

### Participants

Fifty-two students from the University of Trento participated in the study. An additional 4 volunteers were excluded from analysis because they did not complete the task. Participants were aged between 19 and 39 years (32 female, M = 23.25 years, SD = 4.32 years) and were native Italian speakers. None of the participants reported having problems with their sight (normal or corrected-to-normal vision). They had no history of neurological disorders and were not taking any neurological medications. Prior to the experiment, all participants provided written informed consent and received €10 or course credits as compensation for their time. All methods were carried out in accordance with approved guidelines provided by the University of Trento, Human Research Ethics Committee (Protocol 2018-023). The whole procedure was realized in accordance with the Helsinki Declaration.

### Procedure

Participants were welcomed in a dedicated VR lab, gave consent, and completed demographic information. The experiment consisted of two conditions (OR and DD), in which participants were exposed to a series of panoramic videos in VR, with a total duration of ~45 minutes. The order of conditions was counterbalanced across participants. After the head-mounted display (HMD, Oculus Rift) was fitted, participants – comfortably sitting in a chair - started the experiment by being exposed to either the DD or the OR video series. They were encouraged to freely explore the virtual environment by moving their head. The OR video series consisted of 6 panoramic high-definition naturalistic video clips (2732 × 1366 resolution, 20 fps) with a duration of 50 s (Fig. 1a), presented one after the other with no delay in between with a total duration of 5 minutes. All the OR videos represented naturalistic scenes, such as beaches or cascades, and there was a blind spot of approximately 33-degrees located at the bottom of the sphere due to the field of view of the camera. The DD video series was a modified version of the OR videos using the DeepDream^38^. Immediately after the exposure to videos, participants performed the AUT, Stroop tasks, and the ASC questionnaire (Fig. 1b). Those tasks were administered always in the same order via a computer screen, implemented in OpenSesame^64^.

### DeepDream stimuli

DeepDream is a computational procedure that alters images relying on a pre-trained deep convolutional neural network (CNN), a process also referred to as “algorithmic pareidolia”^37^. The algorithm starts by passing an input image *I* with width (*w*) and height (*h*) through the CNN up to a selected layer *A*^*l*^. The objective function 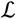 is to maximize the *A*^*l*^ activation. In order to achieve its goal, instead of optimizing the parameters of the network as in the classic approach, it alters the input image by adding the partial derivatives (gradients) of 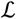 computed with respect to the input image. This optimization algorithm is called gradient ascent because it leads to the maximization of 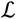. Since DeepDream was conceived for static images, we followed Suzuki et al.^36^ for adapting the algorithm to videos using optical flow to stabilize the optimization process and reduce the variability of the generated frames. In this study, we selected a relative higher layer (inception_4d/pool) of the GoogleNet CNN^38,65^, and setting all the hyperparameters similarly to the Hallucination Machine^36^ (octaves = 3, octave scale = 1.8, iterations = 16, jitter = 32, zoom = 1, step size = 1.5, flow threshold = 6, blending ratio for optical flow = 0.9, blending ratio for background = 0.1).

### The Alternative Uses Task (AUT)

The flexibility of thought was firstly measured by the AUT^39,49^, a widely used divergent thinking task commonly employed for investigating CF^16^. In the AUT, participants were asked to list as many original uses as possible in response to four verbal prompts (“brick”, “newspaper”, “pencil”, “shoe”). They had 2 min to respond to each verbal prompt in a text box, where the responses were constrained to a single or a compound word (e.g. “wrapping paper”, **Fig. 1c**). Participants were instructed to generate single, or compound word responses, instead of the standard open-ended version of the AUT. This allows us to control for the open-ended nature of the task and leads to more consistent and standardized responses by participants. Each condition had two verbal prompts, counterbalanced across participants. As our interest is to study the effect of the DeepDream manipulation on CF, we collapse across AUT items and examine the aggregated action-related space of participants’ responses, and how our manipulation affects this conceptual space.

### Stroop Task

A mouse-tracking version of the Stroop task ^41^ was used to assess CF and the ability to inhibit cognitive interference^66^. The stimuli consisted of the words “red,” “green,” “yellow” and “blue” (in Italian) presented in four response boxes in the upper half of the screen (Fig. 1d). Every trial started after the participant clicked on a start button in the lower right corner of the computer screen, making appear right on top of the start button the target word stimulus after 200ms. Participants were instructed to click on one of the four response boxes corresponding to the ink color of the target stimulus. In 50% of the trials, the color of the displayed word matched its meaning (congruent trials), whereas, in the remaining 50% of the trials, the color and meaning were different (incongruent trials). We also added the constrain that red and green written target words were always printed in either red or green, and blue and yellow always in either blue or yellow. This allowed a fair comparison of the mouse trajectories since the response boxes were placed symmetrically according to this constrain. The total number of trials was 64, congruent and incongruent trials were presented in a randomized order. Mouse trajectories, as well as RT and accuracy, were recorded via the Mousetrap^67^ plugin in Opensesame with a sample rate of 100 Hz (OS: Windows 10, default mouse settings).

### Alternate State of Consciousness questionnaire (ASC)

A short version of the ASC Questionnaire^42^ was administered to control the subjective effects of the videos on certain dimensions of the subjective experience. Indeed, we measured only dimensions that were already found significantly similar to both the Hallucination Machine and the actual psychedelic experience^36^ (Fig. 1e). In the ASC, participants were instructed to rate, on a continuous linear scale (0 = “no more than usual”, 1= “yes, much more than usually”), their experience with the video series as compared to normal waking consciousness.

### Semantic network analysis

Traditionally, using the AUT, most of the research has employed scoring methods of CF based on human judgment^49^ measuring the flexible extent of a participant’s performances. Although this approach has been seen to offer some degree of utility, it still poses some concerns related to the complexities of subjective judgment, raters’ experience, and labor costs. Thus, we opted to quantify CF from the AUT data using two complementary network science approach^4,15^. Following a recently developed^50^ yet extensively applied framework^14^, we modeled the AUT responses as a network in which the nodes represent possible actions (i.e. unique responses) generated by participants in the sample to all of the different AUT cue words, and edges represent relations between two of them. This association indicates the participants’ tendency to generate a word “b” given a word “a” is formed, allowing us to investigate the overall group differences in the network depending on how frequently responses co-occurred across participants^50^.

The raw responses to the AUT, constrained to a single or a compound word, were preprocessed by excluding idiosyncratic answers and non-words and controlling for other possible confounds (i.e., spell-checked, converted plural words into singular). We used McNemar’s chi-squared test to analyze whether there was a difference in proportion between the total number of unique responses and the number of unique responses given by the participants in each condition. In order to construct the cognitive network, we structured the data into a binary *N*×*M* matrix for each condition, in which each column *M* represents the unique response given by all the participants to the AUT verbal prompts, and each row *N* represents a single participant. On each cell, responses were encoded as 1 when the participant *N* provided the response *M* and 0 otherwise. From the binary matrices, we selected only the unique responses (i.e., the columns) generated by at least two participants on each condition and we matched these unique responses between conditions. This allowed us to control for possible confounders, such as the different nodes or edges between conditions^68^. Thus, we constructed a word-similarity matrix by computing the cosine similarity between all the pairs of unique responses for each condition. The resulted matrix is an adjacency matrix of a weighted, fully connected network, having unique word responses as columns and rows and cells as the weight of the link between all the pairs of words. For the sake of retaining the most relevant information in the networks, we removed spurious associations (i.e. weak similarity) by filtering the adjacency matrices with the Triangulated Maximally Filtered Graph (TMFG) method^52^.

The multifaceted aspects of the structure of the cognitive networks was quantified using the following topological quantifiers, after binarizing the networks: CC, ASPL^53^, Q^54^ and S^55^. The CC is an index of how close nodes in a network tend to cluster together and it should be interpreted as a measure of connectivity. Thus, a higher CC implies better local organization and shows stronger connectivity within the network. The ASPL indicates the average number of steps along the shortest paths for all possible pairs of network nodes. A lower value of ASPL might improve the chances of reaching faster relatively remote nodes. The Q assesses how a network is broken down into subnetworks, by quantifying the ways in which a network is divided into sub-networks, while the S can be considered as an index of network flexibility. Indeed, high local connectivity (higher CC) and short global distances between nodes (lower ASPL) define a small-world network, which can be quantified as the ratio between CC and ASPL^53^.

Statistical analysis was conducted by applying two complementary approaches, the LONO and LOSO procedures. In the LONO procedure, we iteratively computed the before mentioned network measures on the partial networks resulting from the exclusion of one node at each iteration, for every node. In the LOSO procedure, in each iteration, we excluded one participant and repeat the pipeline for building the semantic networks and computing the network measures, for every participant. We used a two-tailed paired-samples permutation *t*-test (α = .05, 10000 iterations) to investigate the statistical differences between conditions for each measure in both LONO and LOSO. We also used Cohen’s *d* as a measure of effect size. These analyses were conducted in R using the NetworkToolbox package^69^; inferential statistics and data visualization was implemented in Python using the NetworkX library^70^.

Network percolation analysis estimates the robustness of complex networks under targeted attacks^**71**^. In this study, we implemented percolation analysis^4^ using the weighted TMFG-filtered networks previously constructed from the AUT data. In the percolation analysis, networks are “attacked” by removing links with weight strength below an increasing threshold, called the percolation step ^4^. The initial threshold was the smallest weight in the network and the lowest difference between the sorted weights was used to determine the threshold resolution. In each percolation step, we measure the LCCS, which is the size of the largest connected component, defined as the largest cluster of nodes connected only to each other. The percolation process was terminated when the number of nodes in the largest connected component was less than 3.

Once the percolation process reached its end, we computed *ϕ* (percolation integral), which is the area under the curve representing the LCCS across the percolation steps. It is also formally defined as the sum of all LCCS weighted by their weight threshold value^4^. This measure allowed us to estimate how fast the network breaks apart, a measure of its robustness and structure flexibility. To determine the statistical significance of the percolation analysis results, the LONO, LOSO, and LS were applied as complementary approaches. Similar to the procedures described in the semantic network analysis section, when computing the LONO and LOSO procedures, we iteratively excluded one node or participant, ran the percolation analysis on the resulted networks, and computed *ϕ*, for each node or participant depending on the LONO or LOSO methods employed, respectively. Furthermore, we performed LS analysis in order to control for the possibility that differences between networks may stem from the differences in the link weights. Here, for each network, we randomly selected two pairs of nodes and exchanged them links network. To ensure that the majority of the links are exchanged, this process is repeated 10 times for every link in the networks (1750 shuffles for the OR network, 1730 shuffles for the DD network). This procedure was repeated with 500 iterations, computing *ϕ* on the link-shuffled network at each iteration. We then conducted a paired two-tailed permutation t-test (α = .05, 10000 iterations) between the *ϕ* of the DD and OR conditions for each procedure.

### Drift Diffusion Conflict Modelling

Stroop data were preprocessed by removing outliers both in terms of RT and accuracies. We excluded one participant from the analysis because of the extremely poor performance at one condition (accuracy = 5%), resulting in a sample of 51 participants for subsequent analyses. RT outliers were removed (8.76%) whenever they exceeded ±3 MAD (181 ms) with respect to the median (774 ms) across all trials and participants. Accuracy and RT data across both conditions and congruent and incongruent trials were compared using a paired-samples two-tail permutation t-*t*est (α = .05, 10000 iterations). Then, we fitted the DCM to the RT and accuracy data for each participant and condition separately. The DCM is a computational model suitable for conflict tasks such as the Stroop^45^, modeling the decision-making of participants under the framework of drift-diffusion models^43^. In a drift-diffusion model, the decision process is usually modeled as a Wiener stochastic process *X*_*t*_ as follows:

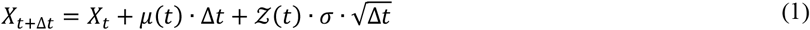

where *μ*(*t*) is the drift of the diffusion process, Δ*t* is the difference between two time points and 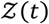 is a random variable that follows a Gaussian distribution 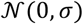 with 0 mean and *σ* standard deviation. In this framework, a decision is made whenever the diffusion process reaches an upper or lower bound *β*. In the DCM, the decision process is modeled as a superimposition of a controlled 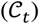 and an automatic 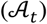 process, as follows:

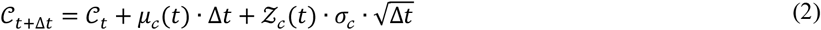

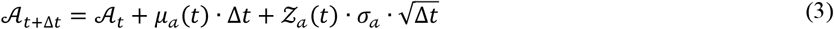

The average time-course of 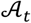 is assumed to follow a rescaled Gamma density function with shape parameter *θ* > 1 and scale parameter *τ*, representing the time-course of the expected mean of 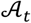. Its amplitude, which is the maximum value, is referred to as *α*, which is positive in congruent trials and negative in incongruent trials. The time-dependent drift *μ*_*a*_ of the automatic process is equal to the first derivative of the expected mean of 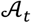 with respect to time *t*, which is:

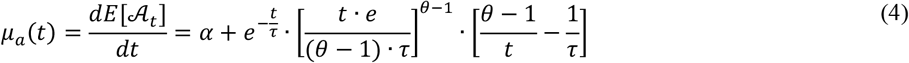

For a more detailed description of the model, see Ulrich et al.^45^. The model was fitted to the RT and accuracy data of each participant and separately for the two conditions, coding the upper bound as the correct response and the lower as the incorrect response. The loss function was the root mean square error (RMSE) between the CDF of the RT and the CAF, which is the proportion of correct responses to targets for different percentiles of the RT distribution, of the simulated data from the DCM against the CDF and CAF of single-subject data. The loss function was minimized using the Nelder-Mead optimizer, with 200 max iterations^72^. We estimated 7 parameters: the amplitude of the automatic process (*α*), the time-to-peak of the automatic process (*τ*), the drift of the controlled process (*δ*), the decision boundary (*β*), the mean and the standard deviation of the non-decisional component and the shape parameter of the starting point. The starting point was kept fixed and the drift rate was constant across trials. We also fixed the standard deviation of the diffusion process to 4 as well as the shape parameter of the automatic process to 2. To explore plausible starting points for the optimization process, the DCM was fitted to each participant’s data using 5000 parameter sets that were randomly generated from a uniform distribution with 5000 trials simulated (see **Table 5** in *Supplementary Materials* for maximum and minimum values). We then took the 15 best parameter sets resulting from this initial search (lowest RMSE) and reran the DCM with 10000 trials 3 times, to avoid local minima^73^. After the process was completed, we took the single best fitting parameter set for each participant and condition. We only analyzed 4 estimated parameters (*α*, *τ*, *δ*, *β*) since they were the ones with a cognitive interpretation that could fit the aims of this study. Finally, we used a two-tailed paired-samples permutation *t*-test (α = .05, 10000 iterations) to investigate the statistical differences between conditions for each selected parameter. Cohen’s *d* was used as a measure of effect size. These analyses were conducted in R using the DCMfun library^74^; inferential statistics and data visualization was implemented in Python.

### Stroop mouse trajectory analysis

Stroop data were also analyzed in terms of mouse trajectories. Trajectories were firstly selected based on the exclusion criteria from RT and accuracy data and including only the correct trials. We extracted the actual motion from the raw trajectories by selecting only the data points in which the cursor actually moved along both the x and y coordinates. Then, we time normalized the trajectories using linear interpolation, making all trajectories with the same number of data points (n = 101). Spatial alignment was performed in order to have the same initial and ending point according to the following equation:

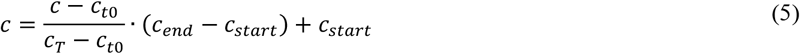

where *c* is either the x or y coordinate time series, *c*_*t0*_ and *c*_*T*_ are the first and last time point and *c*_*start*_ and *c*_*end*_ are the desired initial and ending points. After these preprocessing steps, we computed the AUC between the preprocessed trajectories and the optimal path going from the start button and the target box. We also computed the PE^75^ separately for the x and y coordinates time series, with an embedding dimension of 5 and a time delay of 1. Similar to PE, we computed D as the number of changes in the direction of the cursor along the x and y coordinates. ED was computed as the Euclidean distance between consecutive pairs of coordinates along the whole trajectory, while V represents the ED divided by the time difference. Moreover, we applied a GMM^76^, an unsupervised machine learning algorithm that finds clusters in the data, to estimate macro-states in the mouse trajectories using the scikit-learn library^77^. We ran a series of GMMs in order to select the best number of clusters, using the Expectation-Maximization (EM) algorithm with 1000 maximum iteration and a tolerance criterion of 0.001. The number of clusters varied from 2 to 10 since each trajectory had 101 time points and we wanted to theoretically observe all the transitions between states^78^. Model selection was achieved using the Akaike Information Criterion (AIC) with the constrain that all the clusters had to be present in each participant’s trajectories. The winning GMM was the one with 4 clusters (see supplementary materials, Fig. S1). Then, we computed the transition matrices among conditions by computing the transition probabilities between each pair of states. Finally, we computed the DT as the average lifetime of each state in a trajectory. Statistical significance was assessed with a two-tailed paired-samples permutation t-test (α=0.05, 10000 iterations) and Cohen’s *d* was used as a measure of effect size.

## Author contributions statement

C.R., A.G., and N.D.P. conceived and designed the study. A.G. created the video materials. C.R. and A.G. carried out the experiment, performed the analysis, and lead in writing the manuscript. N.D.P and C.F. assisted with measurements. Y.N.K. C.F. and N.D.P provided critical feedback and helped to shape the research and manuscript. N.D.P supervised the project.

## Additional information

The datasets and video materials used in the current study are available from the corresponding authors on reasonable request.

## Ethics declaration

The authors declare that they have no competing interests.

## References

1. Uddin, L. Q. Cognitive and behavioural flexibility: neural mechanisms and clinical considerations. Nat Rev Neurosci 22, 167–179 (2021).

2. Scott, W. A. Cognitive complexity and cognitive flexibility. Sociometry 405–414 (1962).

3. Mednick, S. The associative basis of the creative process. Psychological review 69, 220–232 (1962).

4. Kenett, Y. N. et al. Flexibility of thought in high creative individuals represented by percolation analysis. Proceedings of the National Academy of Sciences 115, 867–872 (2018).

5. De Pisapia, N. & Rastelli, C. Creativity as an information-based process.

6. Diamond, A. & Lee, K. Interventions shown to aid executive function development in children 4 to 12 years old. Science 333, 959–964 (2011).

7. Burke, S. N. et al. What are the later life contributions to reserve, resilience, and compensation? Neurobiology of aging 83, 140–144 (2019).

8. Waltz, J. A. The neural underpinnings of cognitive flexibility and their disruption in psychotic illness. Neuroscience 345, 203–217 (2017).

9. Kenett, Y. N., Gold, R. & Faust, M. The hyper-modular associative mind: a computational analysis of associative responses of persons with Asperger syndrome. Language and Speech 59, 297–317 (2016).

10. Davidson, M. C., Amso, D., Anderson, L. C. & Diamond, A. Development of cognitive control and executive functions from 4 to 13 years: Evidence from manipulations of memory, inhibition, and task switching. Neuropsychologia 44, 2037–2078 (2006).

11. Edl, S., Benedek, M., Papousek, I., Weiss, E. M. & Fink, A. Creativity and the Stroop interference effect. Personality and individual Differences 69, 38–42 (2014).

12. Logue, S. F. & Gould, T. J. The neural and genetic basis of executive function: attention, cognitive flexibility, and response inhibition. Pharmacology Biochemistry and Behavior 123, 45–54 (2014).

13. Guarino, A., Forte, G., Giovannoli, J. & Casagrande, M. Executive functions in the elderly with mild cognitive impairment: a systematic review on motor and cognitive inhibition, conflict control and cognitive flexibility. Aging & mental health 24, 1028–1045 (2020).

14. Kenett, Y. N. & Faust, M. A Semantic Network Cartography of the Creative Mind. Trends in Cognitive Sciences 23, 271–274 (2019).

15. Kenett, Y. N., Anaki, D. & Faust, M. Investigating the structure of semantic networks in low and high creative persons. Frontiers in human neuroscience 8, 407 (2014).

16. Ritter, S. M. et al. Diversifying experiences enhance cognitive flexibility. Journal of experimental social psychology 48, 961–964 (2012).

17. Chirico, A., Glaveanu, V. P., Cipresso, P., Riva, G. & Gaggioli, A. Awe enhances creative thinking: an experimental study. Creativity Research Journal 30, 123–131 (2018).

18. Glass, B. D., Maddox, W. T. & Love, B. C. Real-time strategy game training: emergence of a cognitive flexibility trait. PloS one 8, e70350 (2013).

19. Ritter, S. M. & Mostert, N. Enhancement of creative thinking skills using a cognitive-based creativity training. Journal of Cognitive Enhancement 1, 243–253 (2017).

20. Baggott, M. J. Psychedelics and creativity: a review of the quantitative literature. PeerJ PrePrints 3, e1202v1 (2015).

21. Family, N. et al. Semantic activation in LSD: evidence from picture naming. Language, Cognition and Neuroscience 31, 1320–1327 (2016).

22. Prochazkova, L. et al. Exploring the effect of microdosing psychedelics on creativity in an open-label natural setting. Psychopharmacology 235, 3401–3413 (2018).

23. Murphy-Beiner, A. & Soar, K. Ayahuasca’s ‘afterglow’: improved mindfulness and cognitive flexibility in ayahuasca drinkers. Psychopharmacology 237, 1161–1169 (2020).

24. Mason, N. L. et al. Spontaneous and deliberate creative cognition during and after psilocybin exposure. Translational psychiatry 11, 1–13 (2021).

25. Girn, M., Mills, C., Roseman, L., Carhart-Harris, R. L. & Christoff, K. Updating the dynamic framework of thought: Creativity and psychedelics. Neuroimage 213, 116726 (2020).

26. Uthaug, M. et al. Sub-acute and long-term effects of ayahuasca on affect and cognitive thinking style and their association with ego dissolution. Psychopharmacology 235, 2979–2989 (2018).

27. Kuypers, K. P. C. et al. Ayahuasca enhances creative divergent thinking while decreasing conventional convergent thinking. Psychopharmacology 233, 3395–3403 (2016).

28. Sessa, B. Is it time to revisit the role of psychedelic drugs in enhancing human creativity? Journal of Psychopharmacology 22, 821–827 (2008).

29. Harman, W. W., McKim, R. H., Mogar, R. E., Fadiman, J. & Stolaroff, M. J. Psychedelic agents in creative problem-solving: A pilot study. Psychological reports 19, 211–227 (1966).

30. McGlothlin, W., Cohen, S. & McGlothlin, M. S. Long lasting effects of LSD on normals. Archives of General Psychiatry 17, 521–532 (1967).

31. Carhart-Harris, R. L. et al. Neural correlates of the LSD experience revealed by multimodal neuroimaging. Proceedings of the National Academy of Sciences 113, 4853–4858 (2016).

32. Carhart-Harris, R. L. & Friston, K. REBUS and the anarchic brain: toward a unified model of the brain action of psychedelics. Pharmacological reviews 71, 316–344 (2019).

33. Preller, K. H. et al. The fabric of meaning and subjective effects in LSD-induced states depend on serotonin 2A receptor activation. Current Biology 27, 451–457 (2017).

34. Spitzer, M. et al. Increased activation of indirect semantic associations under psilocybin. Biological psychiatry 39, 1055–1057 (1996).

35. Carhart-Harris, R. L. et al. The entropic brain: a theory of conscious states informed by neuroimaging research with psychedelic drugs. Frontiers in Human Neuroscience 8, 20 (2014).

36. Suzuki, K., Roseboom, W., Schwartzman, D. J. & Seth, A. K. A deep-dream virtual reality platform for studying altered perceptual phenomenology. Scientific Reports 7, 1–11 (2017).

37. Greco, A., Gallitto, G., D’Alessandro, M. & Rastelli, C. Increased Entropic Brain Dynamics during DeepDream-Induced Altered Perceptual Phenomenology. Entropy 23, 839 (2021).

38. Mordvintsev, A., Olah, C. & Tyka, M. Inceptionism: Going Deeper into Neural Networks. Google Research Blog Available at: http://googleresearch.blogspot.co.uk/2015/06/inceptionism-going-deeper-into-neural.html. http://googleresearch.blogspot.co.uk/2015/06/inceptionism-going-deeper-into-neural.html (2015).

39. Guilford, J. P. The nature of human intelligence. (1967).

40. Stroop, J. R. Studies of interference in serial verbal reactions. Journal of experimental psychology 18, 643 (1935).

41. Bundt, C., Ruitenberg, M. F., Abrahamse, E. L. & Notebaert, W. Early and late indications of item-specific control in a Stroop mouse tracking study. PloS one 13, e0197278 (2018).

42. Dittrich, A. The standardized psychometric assessment of altered states of consciousness (ASCs) in humans. Pharmacopsychiatry 31, 80–84 (1998).

43. Ratcliff, R. A theory of memory retrieval. Psychological review 85, 59 (1978).

44. Ratcliff, R. & McKoon, G. The diffusion decision model: theory and data for two-choice decision tasks. Neural computation 20, 873–922 (2008).

45. Ulrich, R., Schröter, H., Leuthold, H. & Birngruber, T. Automatic and controlled stimulus processing in conflict tasks: Superimposed diffusion processes and delta functions. Cognitive psychology 78, 148–174 (2015).

46. Cosgrove, A. L., Kenett, Y. N., Beaty, R. E. & Diaz, M. T. Quantifying flexibility in thought: The resiliency of semantic networks differs across the lifespan. Cognition 211, 104631 (2021).

47. Rastelli, C., Greco, A. & Finocchiaro, C. Revealing the Role of Divergent Thinking and Fluid Intelligence in Children’s Semantic Memory Organization. J. Intell. 8, 43 (2020).

48. Sanders, J., Johnson, K. A., Garavan, H., Gill, M. & Gallagher, L. A review of neuropsychological and neuroimaging research in autistic spectrum disorders: Attention, inhibition and cognitive flexibility. Research in autism spectrum disorders 2, 1–16 (2008).

49. Torrance, E. P. Norms-technical manual: Torrance Tests of Creative Thinking Lexington. MA: Ginn & Co (1974). 19

50. Kenett, Y. et al. Semantic organization in children with cochlear implants: computational analysis of verbal fluency. Frontiers in Psychology 4, 543 (2013).

51. Collins, A. M. & Loftus, E. F. A spreading-activation theory of semantic processing. Psychological Review 82, 407–428 (1975).

52. Massara, G. P., Di Matteo, T. & Aste, T. Network filtering for big data: Triangulated maximally filtered graph. Journal of complex Networks 5, 161–178 (2017).

53. Watts, D. J. & Strogatz, S. H. Collective dynamics of ‘small-world’ networks. Nature 393, 440–442 (1998).

54. Newman, M. E. J. Modularity and community structure in networks. PNAS 103, 8577–8582 (2006).

55. Humphries, M. D. & Gurney, K. Network ‘Small-World-Ness’: A Quantitative Method for Determining Canonical Network Equivalence. PLOS ONE 3, e0002051 (2008).

56. Siew, C. S., Wulff, D. U., Beckage, N. M. & Kenett, Y. N. Cognitive network science: A review of research on cognition through the lens of network representations, processes, and dynamics. Complexity 2019, (2019).

57. Benedek, M. et al. How semantic memory structure and intelligence contribute to creative thought: a network science approach. Thinking & Reasoning 23, 158–183 (2017).

58. Anderson, J. R. A spreading activation theory of memory. Journal of verbal learning and verbal behavior 22, 261–295 (1983).

59. Kenett, Y. & Thompson-Schill, S. L. Novel conceptual combination can dynamically reconfigure semantic memory networks. (2020).

60. Kumar, A. A. Semantic memory: A review of methods, models, and current challenges. Psychonomic Bulletin & Review 28, 40–80 (2021).

61. Yee, E. & Thompson-Schill, S. L. Putting concepts into context. Psychonomic bulletin & review 23, 1015–1027 (2016).

62. Gelfo, F. Does experience enhance cognitive flexibility? An overview of the evidence provided by the environmental enrichment studies. Frontiers in behavioral neuroscience 13, 150 (2019).

63. Yan, T. et al. Left temporal pole contributes to creative thinking via an individual semantic network. Psychophysiology e13841 (2021).

64. Mathôt, S., Schreij, D. & Theeuwes, J. OpenSesame: An open-source, graphical experiment builder for the social sciences. Behavior research methods 44, 314–324 (2012).

65. Szegedy, C. et al. Going deeper with convolutions. in Proceedings of the IEEE conference on computer vision and pattern recognition 1–9 (2015).

66. Scarpina, F. & Tagini, S. The stroop color and word test. Frontiers in psychology 8, 557 (2017).

67. Kieslich, P. J. & Henninger, F. Mousetrap: An integrated, open-source mouse-tracking package. Behav Res 49, 1652–1667 (2017).

68. Van Wijk, B. C., Stam, C. J. & Daffertshofer, A. Comparing brain networks of different size and connectivity density using graph theory. PloS one 5, e13701 (2010).

69. Christensen, A. P. NetworkToolbox: Methods and Measures for Brain, Cognitive, and Psychometric Network Analysis in R. R J. 10, 422 (2018).

70. Hagberg, A., Swart, P. & S Chult, D. Exploring network structure, dynamics, and function using NetworkX. (2008).

71. CohenR, H. 2010Complex networks: structure, robustness and function.

72. Nelder, J. A. & Mead, R. A simplex method for function minimization. The computer journal 7, 308–313 (1965).

73. Hedge, C., Vivian-Griffiths, S., Powell, G., Bompas, A. & Sumner, P. Slow and steady? Strategic adjustments in response caution are moderately reliable and correlate across tasks. Consciousness and Cognition 75, 102797 (2019).

74. Mackenzie, I. G. & Dudschig, C. DMCfun: An R package for fitting Diffusion Model of Conflict (DMC) to reaction time and error rate data. Methods in Psychology 5, 100074 (2021).

75. Bandt, C. & Pompe, B. Permutation Entropy: A Natural Complexity Measure for Time Series. Phys. Rev.Lett. 88, 174102 (2002).

76. Reynolds, D. A. Gaussian mixture models. Encyclopedia of biometrics 741, 659–663 (2009).

77. Pedregosa, F. et al. Scikit-learn: Machine learning in Python. the Journal of machine Learning research 12, 2825–2830 (2011).

78. Cornblath, E. J. et al. Temporal sequences of brain activity at rest are constrained by white matter structure and modulated by cognitive demands. Communications biology 3, 1–12 (2020).

