## Supplementary Materials for "Simulated visual hallucinations in virtual reality enhance cognitive flexibility"

### 1. ASC questionnaire results

| Dimension | Item | OR | DD | $t\ (p)$ | $d$ |
|---|---|---|---|---|---|
| Arousal | "Please rate your general level of emotional arousal during the last session" | 0.57 (0.26) | 0.57 (0.27) | 0.209 (0.834) | 0.03 |
| Ego | "I experienced a dissolving of my 'self' or 'ego'" | 0.44 (0.28) | 0.44 (0.26) | 0.032 (0.974) | 0.01 |
| Imagey | "I saw complex visual imagery" | 0.30 (0.25) | 0.88 (0.19) | -12.832 (<0.001) | 2.63 |
| Intensity | "Please rate the intensity of the experience during the last session" | 0.52 (0.24) | 0.63 (0.25) | -2.894 (0.01) | 0.46 |
| Muddle | "My think was muddled" | 0.32 (0.22) | 0.60 (0.32) | -5.073 (<0.001) | 1.03 |
| Patterns | "I saw patterns and colours" | 0.45 (0.26) | 0.86 (0.19) | -9.459 (<0.001) | 1.79 |
| Space | "My sense of size and space was distorted" | 0.39 (0.27) | 0.64 (0.33) | -4.293 (<0.001) | 0.81 |
| Spirit | "The experience had a spiritual or mystical quality" | 0.47 (0.25) | 0.60 (0.3) | -2.514 (0.01) | 0.50 |
| Strange | "Things looked strange" | 0.39 (0.2) | 0.92 (0.18) | -15.949 (<0.001) | 2.85 |
| Vivid | "My imagination was extremely vivid" | 0.44 (0.29) | 0.57 (0.28) | -2.88 (0.015) | 0.45 |

**Table 1. Results of the comparison between OR and DD condition on ASC.** Mean and standard deviation in parenthesis. Results from two-tailed paired-samples permutation t-test with 10000 permutations. The t-statistics is calculated from a parametric t-test student; a negative t-statistics denote the OR condition having lower values that the DD condition. Choen's d effect sizes: < 0.20, negligible; 0.20, small; 0.50, moderate; 0.80, large; 1.10, very large.

### 2. Minimum and maximum values of uniform distributions used to generate starting parameters for the DCM

| | Minimum | Maximum |
|---|---|---|
| Amplitude ($\alpha$) | 15 | 40 |
| Time-to-peak ($\tau$) | 100 | 600 |
| Drift rate ($\delta$) | 2 | 8 |
| Boundary ($\beta$) | 30 | 80 |
| Non-decision mean | 270 | 400 |
| Non-decision standard deviation | 20 | 50 |
| Start shape | 1 | 10 |
